# Expanded Asgard archaea shed new light on the origin of eukaryotes and support a 2-domain tree of life

**DOI:** 10.1101/2021.02.04.429862

**Authors:** Ruize Xie, Yinzhao Wang, Danyue Huang, Jialin Hou, Liuyang Li, Haining Hu, Xiaoxiao Zhao, Fengping Wang

## Abstract

The hypothesis that eukaryotes originated from within the domain Archaea has been strongly supported by recent phylogenomic analyses placing Heimdallarchaeota from the Asgard superphylum as the closest known archaeal sister-group to eukaryotes. At present, only seven phyla are described in the Asgard superphylum, which limits our understanding of the relationship between eukaryotes and archaea, as well as the evolution and ecological functions of Asgard archaea. Here, we describe five novel phylum-level Asgard archaeal lineages, tentatively named Tyr-, Sigyn-, Freyr-, Hoder- and Balderarchaeota. Comprehensive phylogenomic analyses supported a new Asgard lineage Tyrarchaeota was identified as a deeper branching lineage cluster with the eukaryotic nuclear host lineage than Heimdallarchaeota that were previously considered as the closest archaeal relatives of eukaryotes. Metabolic reconstruction of Tyrarchaeota suggests a mixotrophic lifestyle of this archaea, capable of peptides and amino acids utilization while having the potential using the Wood-Ljungdahl pathway for carbon fixation and acetogenesis. This study largely expands the Asgard superphylum, provides additional evidences to support the 2-domain life tree thus sheds new light on the evolution and geochemical functions of the Asgard archaea.

## Introduction

The origin of eukaryotes is considered as a critical biological evolutionary event on Earth [1–4]. The common ancestor of eukaryotes is generally believed to have evolved from a symbiotic process [5,6] in which one endosymbiotic bacterium within the Proteobacteria phylum evolved into a mitochondrion [7,8] and one endosymbiotic host cell became the cell nucleus [9–11]. The identity of the host cell ancestor has been vigorously debated, and two hypotheses regarding 2- or 3-domain trees of life have been raised [12,13]. However, increasing evidence provided by phylogenomic analyses [10,14], as well as the presence of eukaryotic signature proteins (ESPs) [15] in the Asgard archaea, has supported the idea that eukaryotic cells originated in the domain Archaea, particularly in the archaeal Asgard superphylum [9,10,16]. The Asgard archaea are described as mixotrophic or heterotrophic [11,17] and are ubiquitously distributed in various environments, such as hydrothermal vents [9,10]; lake, river and marine sediments [18]; microbial mats [19]; and mangroves [17]. These organisms potentially play important roles in global element cycling [20]. The identification of Lokiarchaeota in the Loki’s Castle hydrothermal vent field provided pivotal genomic and phylogenetic evidence that eukaryotes originated within the domain Archaea, supporting a 2-domain tree of life, which is consistent with the eocyte hypothesis [9]. Further discovery and proposal of the Asgard superphylum have provided new insights into the transition of archaea to eukaryotes and into the origin of eukaryotic cell complexity [10]. Within the Asgard superphylum, Heimdallarchaeota has been identified to be the closest Asgard archaeal lineage to the eukaryotic branch on the phylogenetic tree on the basis of carefully selected conserved protein sequences [10,14]. Recently, Imachi *et al.* cultivated one Asgard archaeon, *Candidatus* Prometheoarchaeum syntrophicum strain MK-D1, in the laboratory and observed, for the first time, the intertwining of this archaeon with bacterial cells via extracellular protrusions under a transmission electron microscope [21]. The idea of an archaeal origin of eukaryotes and a 2-domain tree of life has recently become increasingly favorable [14,22]; nevertheless, our understanding of the evolution of Asgard archaea, the archaea-eukaryote transition, and the geochemical roles of these evolutionarily important archaea remains incomplete. This lack of understanding is largely due to the limited number of high-quality genomes of Asgard archaea, which are considered highly diverse as revealed by 16S rRNA gene surveys [20,23]; yet only a small fraction have representative genomes. In this study, we assembled five previously unknown phylum-level Asgard archaeal group, greatly expanded the Asgard genomic diversity.

## Results

### Expanded Asgard archaea support 2-domain tree of life

Raw metagenomic data from various environments were downloaded from the publicly available National Center for Biotechnology Information (NCBI) Sequence Read Archive (SRA) database (Supplementary Table 1). The data were subsequently reassembled, binned and classified as described in the Methods section. In total, 128 Asgard MAGs were obtained and in-depth phylogenomic analyses were performed on 37 concatenated conserved proteins [24] under LG+R10 model to confirm the placement of these MAGs on phylogenomic tree (Supplementary Table 2). The analysis revealed that, in addition to the previously described Loki-, Thor-, Odin-, Heimdall-, Hela-, Gerd- and Hermodarchaeota clades [25], there are five additional monophyletic branching clades (Fig. 1 and Supplementary Fig. 3), here tentatively named Tyr-, Sigyn-, Freyr-, Hoder- and Balderarchaeota after the Asgard gods in the Norse mythology (Tyr, the god of war; Sigyn, a goddess of victory; Freyr, the god of peace; Hoder, the god of darkness; and Balder, the god of light). The MAGs of these new Asgard lineages were recovered from different environments (Tyrarchaeota was found in coastal sediments, Sigynarchaeota and Hoderarchaeota were derived from lake sediments, Freyrarchaeota was identified in marine sediments and Balderarchaeota was detected in marine and freshwater sediments). Near-complete MAGs ranging in size from 2.4 to 6.2 Mb with completeness ranging from 90.65 to 96.26% were constructed for representatives of each new Asgard clade (Supplementary Table 3). To further assess their distinctiveness compared to the Asgard members already defined, we calculated the average nucleotide identity (ANI) (Supplementary Fig. 1) and average amino acid identity (AAI) (Supplementary Fig. 2) between them and other Asgard MAGs. The AAI values showed that all the MAGs discovered here share a low AAI with the known Asgard archaea (<50%) and fall within the phylum-level classification range (40%~52%) [26], providing additional support for the uniqueness of these new Asgard lineages.

**Figure 1.**
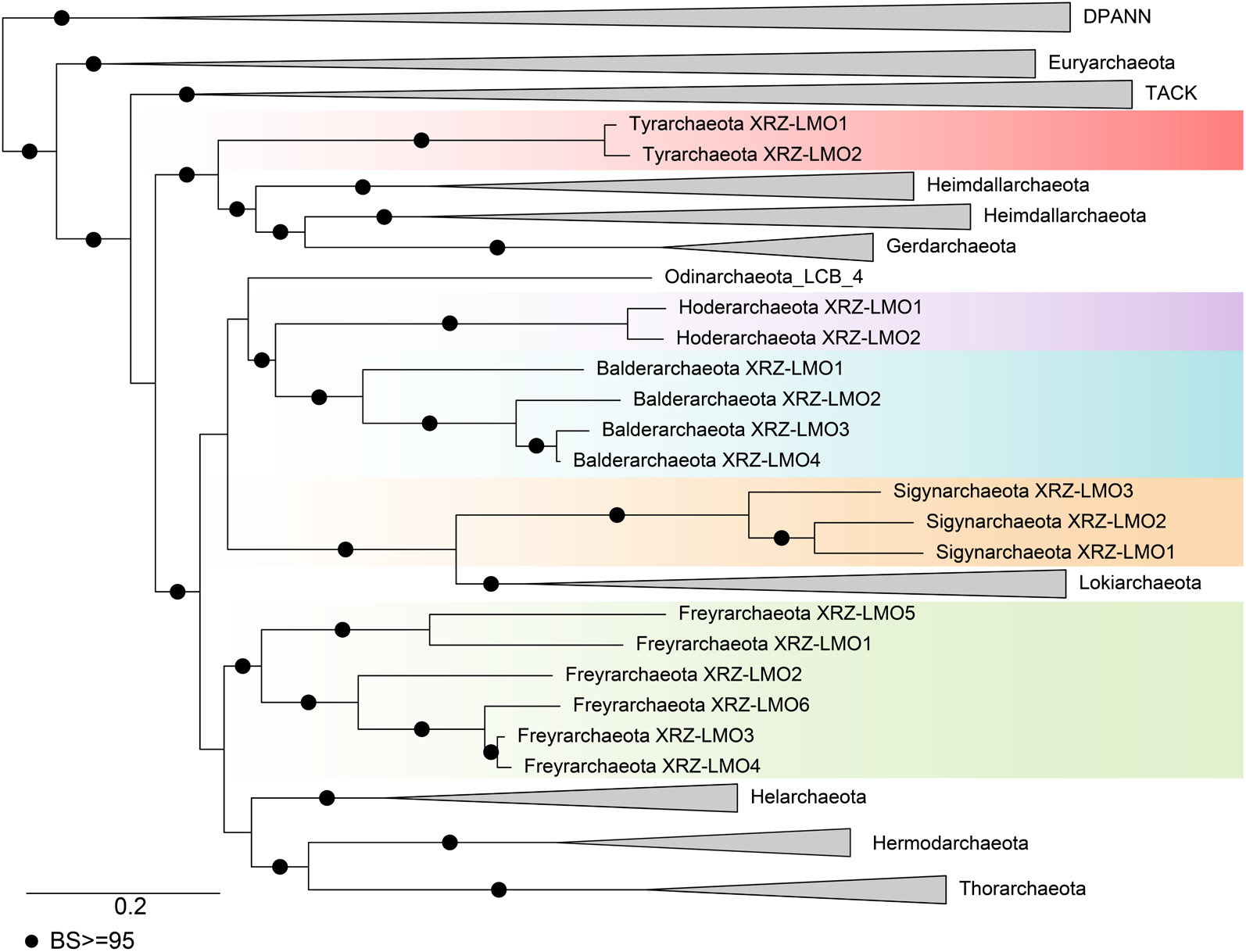
Phylogenetic tree of the Asgard archaea using DPANN as an outgroup. Maximum-likelihood tree of 37 concatenated marker proteins inferred with the LG+R10 model in IQ-TREE; The bootstrap support values above 95 were shown with black filled circles. 19 representatives of DPANN, 55 representatives of Euryarchaeota, 29 representatives of TACK and 68 genomes (including five new lineages) of Asgard were used to infer phylogenetic tree. The initial tree is at supplementary Fig. 3.

To determine the phylogenetic positions of these new Asgard lineages in relation to eukaryotes, we performed maximum-likelihood and Bayesian inference phylogenetic analyses using 21 conserved marker proteins carefully selected by Williams *et al*. [14] and maximum likelihood phylogenetic analysis of 55 archaeal-eukaryotic ribosomal proteins [10]. The taxa included in these analyses were also selected on the basis of the instructions of Williams *et al.* [14]: a representative taxon set was constructed comprising 74 archaeal genomes (43 within Asgard), 19 eukaryotic genomes, and 36 bacterial genomes (Supplementary Table 4). To avoid potential phylogenetic artifacts resulting from eukaryotic genes from the mitochondria or plastids, single-gene datasets were carefully inspected with single protein trees, and BLASTp inspection was performed. We concatenated two gene sets, then inferred a Bayesian tree under the CAT+GTR+G4 model of 21 marker genes and a maximum likelihood tree of 55 ribosomal proteins under the LG+C60+F+G4 model. Both trees robustly indicate that the eukaryotic lineage branches within the Asgard superphylum with high statistical supports (posterior probability (PP)=0.99, Fig. 2a and Supplementary Fig. 4; bootstrap support (BS) = 90, Fig. 2b and Supplementary Fig. 5). Moreover, both the Bayesian and maximum-likelihood (LG+C60+F+G4 model) phylogenetic analyses placed Tyrarchaeota as the deeper branching clade with eukaryotes than Heimdallarchaeota which was previously described as the sister lineage to eukaryotes [14]. The eukaryotes phylogenetic lineage together with Tyr-, Heimdall- and Gerdarchaeota now form a robust sister clade with the rest of the Asgard archaeal lineages. Considering that inter-archaeal horizontal gene transfers (HGTs) may exist in nonribosomal conserved proteins, which probably distort topology of tree, we filtered out some archaeal genes which acquired by inter-archaeal HGT, then performed additional maximum likelihood phylogenetic analyses using LG+C60+F+G4 model. Consistently, phylogenetic trees of 21 marker genes after removing inter-archaeal HGT genes showed that eukaryotes still form a monophyletic lineage with Tyr-, Heimdall- and Gerdarchaeota (BS=94, Supplementary Fig. 6). In summary, the phylogenomic analyses conducted in this study provide strong support for a 2-domain tree of life in which eukaryotes emerged from within the Asgard clade.

**Figure 2.**
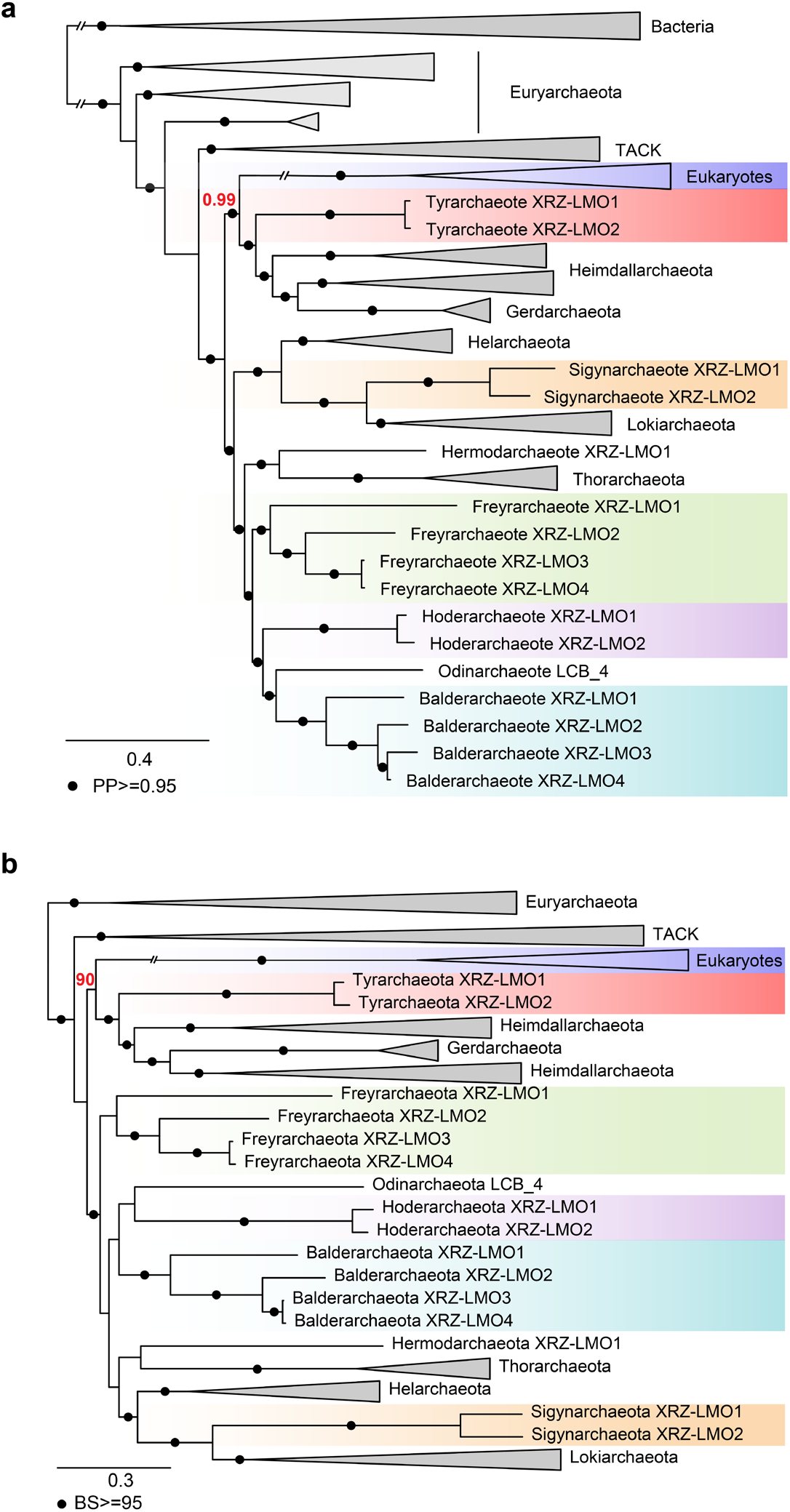
Phylogenetic affiliations of bacteria, archaea and eukaryotes. **a**, Bayesian inference of 21 concatenated conserved protein sequences under the CAT+GTR model rooted in bacteria; **b**, Maximum-likelihood analysis of 55 archaeal-eukaryotic ribosomal proteins under the LG+C60+F+G4 model rooted in Euryarchaeota. Both trees show high support value (PP=0.99; BS=90) that eukaryotes are the sister group of a clade consisting of Tyr-, Heimdall- and Gerdarchaeota. The initial trees are at supplementary Fig. 4-5.

### ESP-encoding genes widely shared by Asgard archaea

The potential ESPs were identified from the newly reconstructed Asgard MAGs (Fig. 3). Consistent with previous reports [9,10,16,27], different key subunits of informational processing machinery were found. For example, topoisomerase IB protein-encoding genes were identified in Freyr-, Balder- and Hoderarchaeota, while Balderarchaeote XRZ-LMO2, XRZ-LMO3, XRZ-LMO4 and Freyrarchaeote XRZ-LMO2 were found to encode a RNA polymerase subunit G. The Balderarchaeota and Tyrarchaeota clades were also found to contain genes related to cell division and the cytoskeleton, and all the newly assembled Asgard archaea share genes coding for gelsolin domain-containing proteins. With regard to actin-related proteins, two to three related subunits were detected in Tyra- and Balderarchaeota, whereas profilin domain protein-encoding genes were identified in Tyr-, Sigyn- and Balderarchaeota.

**Figure 3.**
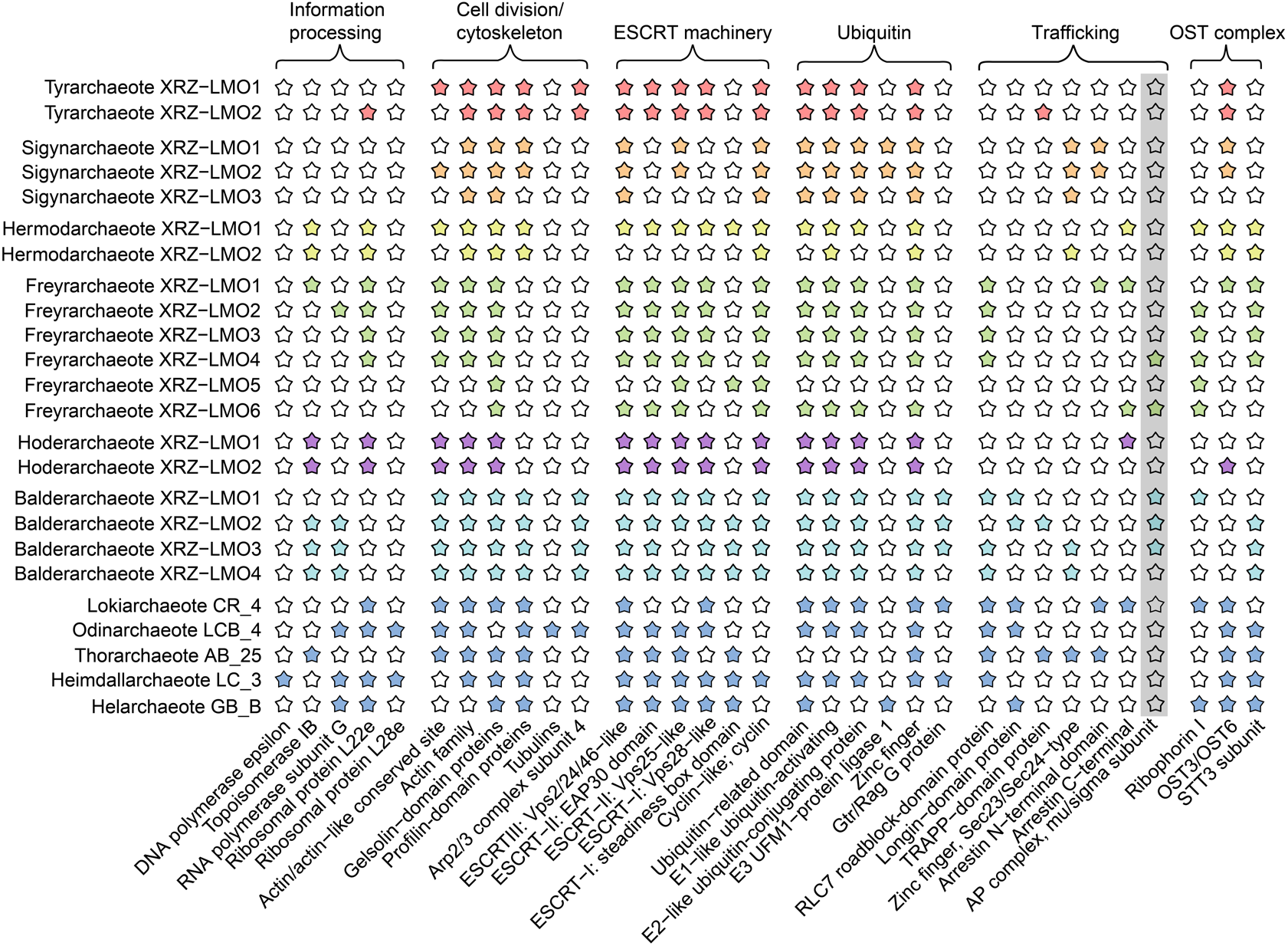
Comparison of the distributions of ESPs in the five new Asgard lineages and other representative Asgard clades. Colored stars indicate the presence of ESPs, whereas empty stars indicate the absence of ESPs. The grep box highlights ESP identified in this study. The new ESP, mu/sigma subunit of AP complex, was detected in Balderarchaeota XRZ-LMO1, Balderarchaeota XRZ-LMO2, Balderarchaeota XRZ-LMO3, Freyrarchaeota XRZ-LMO4 and Freyrarchaeota XRZ-LMO6.

Ubiquitin-based signaling is an important cellular process in eukaryotes [28]. Previous studies have reported the presence of the related protein domains in Loki-, Odin-, Hel- and Heimdallarchaeota but not in Thorarchaeota [9,27,29]. Here, we identified ubiquitin system-related protein-encoding genes in nearly all newly assembled MAGs including several ubiquitin-related domains, zinc fingers, ubiquitin-activating enzyme (E1), ubiquitin-conjugating protein (E2), and UFM1-protein ligase 1 (E3), indicating that the ubiquitin system is widespread in Asgard archaea.

The endosomal sorting complex required for transport (ESCRT) machinery consisting of complexes I-III and associated subunits [9,30,31] were identified in the newly recovered MAGs. Genes coding for Vps28 domain-containing proteins previously found in Loki-, Odin-, Hel- and Heimdallarchaeota [25,27] were also identified in Tyr-, Freyr-, Balder- and Hoderarchaeota but were absent from Sigynarchaeota. The Sigynarchaeota MAGs also lack genes for both EAP30 domain- and steadiness box domain-containing proteins. Notably, all new Asgard MAGs contain cyclin-like protein-encoding genes, whereas Freyrarchaeote XRZ-LMO2 lacks genes for ESCRT complexes I-III.

ESPs with intracellular trafficking and secretion functions were also identified in MAGs here. However, only Balderarchaeota MAGs encode proteins with homology to Gtr/Rag-family GTPases and contain longin domain-encoding genes. All Sigynarchaeota MAGs possess genes coding for zinc finger- and Sec23/24-type proteins. Genes coding for both the N- and C-termini of arrestins were found in Freyrarchaeote XRZ-LMO1; the organization resembles one previously reported in the genome of Lokiarchaeote CR-4, in which the C- and N-terminal domain proteins are separated from each other by one gene [25].

We also analyzed the oligosaccharyltransferase (OST) complex in the reconstructed MAGs, and the results showed that OST complex-encoding genes were present in the MAGs of all five Asgard clades except for Sigynarchaeote XRZ-LMO3 and Hoderarchaeote XRZ-LMO1. Ribophorin I homolog-encoding genes were found in the Hermod- and Balderarchaeota MAGs. Homologs of OST3/6, which have been demonstrated to influence yeast glycosylation efficiency [32], were also identified in several Asgard MAGs, while STT3 subunit protein-encoding genes were detected in the Freyr- and Balderarchaeota MAGs, consistent with previous reports on Loki-, Odin-, Thor-, Hel- and Heimdallarchaeota [25].

In addition to reported ESPs, we identified a potential ESP belonging to mu/sigma subunit of AP (adaptor protein) complex-encoding genes in Balder- and Freyrarchaeota MAGs (Fig. 3). Homologues of mu/sigma subunit of AP complex contain IPR022775 domain. AP complexes are classified into AP-1, AP-2, AP-3, AP-4 and AP-5 and all AP complexes are heterotetramers consisting of two large subunits (adaptins), one medium-sized subunit (mu) and one small-sized subunit (sigma) [33,34]. AP complexes play a vital role in mediating intracellular membrane trafficking [35]. Taken together, the identification of potential ESP provides further insight into the origins of eukaryotic cellular complexity.

### Metabolic reconstructions of newly discovered Asgard lineages

For carbon metabolism, Balder-, Freyr- and Sigynarchaeota contain genes for complete steps of glycolysis via Embden-Meyerhof-Parnas (EMP) pathway and most steps of the tricarboxylic acid (TCA) cycle (Fig. 4). Among genes associated with glycolysis, the specific enzymes catalyzing these steps is difference in three lineages. In the first step of glycolysis, for example, Freyrarchaeota and Sigynarchaeota prone to use ATP-dependent ROK (repressor, open reading frame, kinase) family enzymes but ADP-dependent glucokinase performs this step in Balderarchaeota. Likewise, Freyrarchaeota and Sigynarchaeota encode ATP-dependent phosphofructokinase (PfkB) but fructose 6-phosphate (F6P) to fructose 1,6-bisphosphate (F1,6P) was catalyzed by ADP-dependent phosphofructokinase (ADP-PFK) in Balderarchaeota. In particular, Sigynarchaeote XRZ-LMO2 contains not only all genes responsible for glycolysis and TCA but also a large proportion of genes encoding extracellular carbohydrate-degrading enzymes, including α-amylase, cellulase, α-mannosidases and β-glucosidases (Supplementary Table 5), indicating that archaea in Sigynarchaeota have the capacity to use complex carbohydrates. However, Hoder- and Tyrarchaeota lack some key enzymes for glycolysis and TCA, including hexo- or glucokinase, citrate synthase, and malate dehydrogenase, suggesting that they lack the ability to oxidize carbohydrates. Intriguingly, all five newly assembled Asgard archaeal lineages possess genes in the tetrahydromethanopterin Wood-Ljungdahl (THMPT-WL) pathway (with THMPT as the C1 carrier), while Sigynarchaeota even harbors an additional tetrahydrofolate Wood-Ljungdahl (THF-WL) pathway (with THF as the C1 carrier) (Fig. 4). Both pathways can be used to reduce CO_2_ for acetyl-CoA production, with archaea normally utilizing the THMPT-WL pathway and acetogenic bacteria normally utilizing the THF-WL pathway [36]. In Hoder- and Tyrarchaeota MAGs, glycolysis and TCA pathways are not complete, however, the presence of THMPT-WL pathway genes and group 3 [NiFe] hydrogenases encoding genes implies their potential to harness energy from H_2_ oxidation, possibly for lithoautotrophic growth or utilization of amino acids, depending on environmental condition, as suggested for Lokiarchaeota and Thorarchaeota [37,38]. Moreover, *de novo* anaerobic cobalamin (vitamin B_12_) biosynthesis pathway was found in Tyrarchaeota (Fig. 4), suggesting that Tyrarchaeota harbor the potential of cobalamin synthesis. In nature, only limited members of bacteria and archaea possess capacity of *de novo* cobalamin synthesis using one of two alternative pathways: aerobic or anaerobic pathway [39]. Within Archaea, the Phyla Euryarchaeota, Thaumarchaeota, Crenarchaeota and Bathyarchaeota have been reported possessing cobalamin synthesizing pathway [40,41], while being firstly identified in the Asgard archaea here.

**Figure 4.**
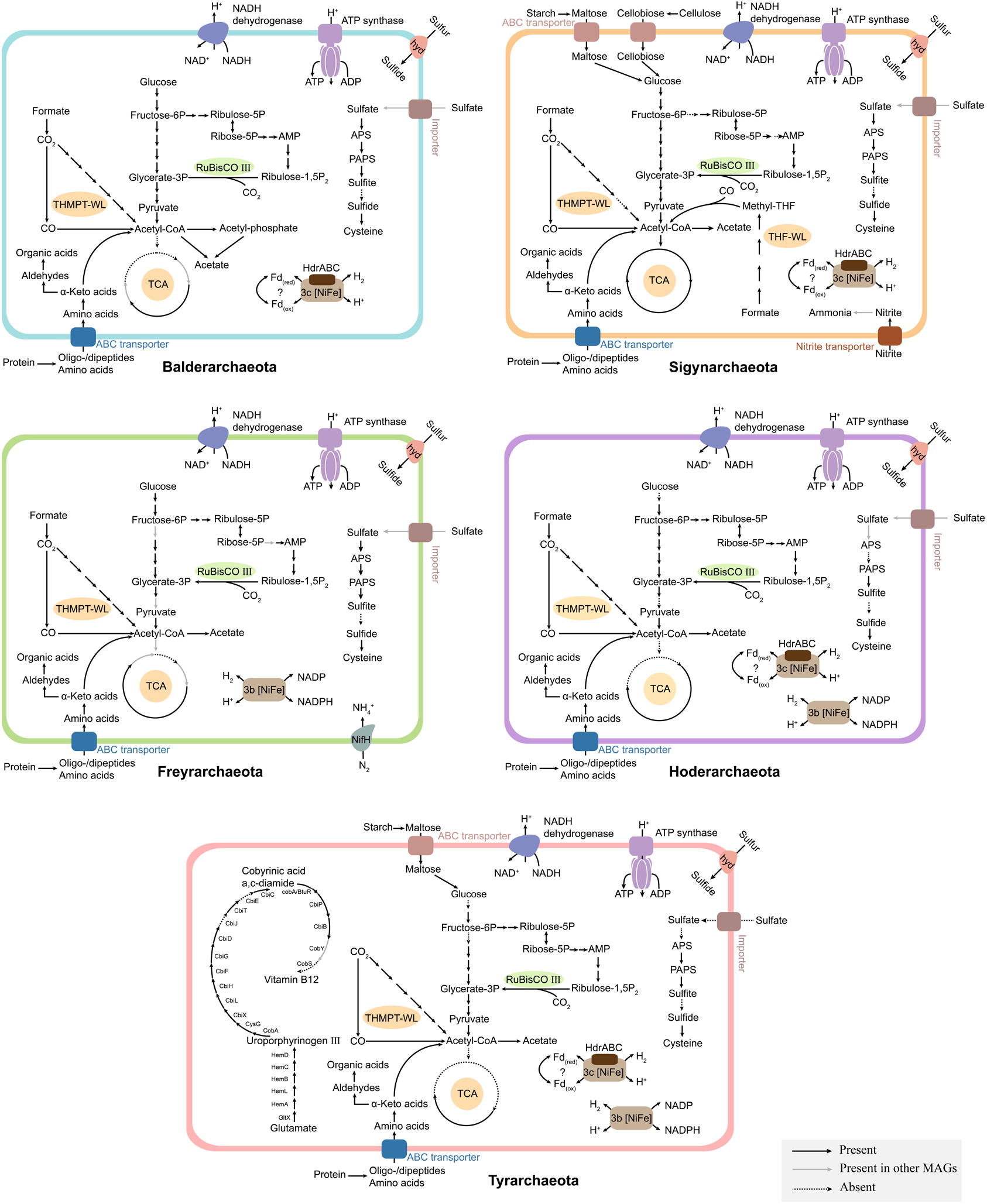
Inferred metabolic pathways of the five new Asgard lineages based on genes identified using the KEGG database and the NCBI NR protein database. A black line indicates that a component/process is present in representative MAGs, a grey line indicates that a component/process is present in other MAGs, and a dashed line indicates that a certain pathway or enzyme is absent from all genomes. The representatives of the different lineages are as follows: Balderarchaeota, XRZ-LMO2; Freyrarchaeota, XRZ-LMO2; Hoderarchaeota, XRZ-LMO1; Sigynarchaeota, XRZ-LMO2; and Tyrarchaeota, XRZ-LMO1. Details about the genes are provided in Supplementary Table 6.

The ADP-dependent acetyl-CoA synthetase (ACD) for acetogenesis, which is widely found in archaea [42], was identified in all newly discovered Asgard lineages. Meanwhile, phosphate acetyltransferase (Pta) and acetate kinase (Ack) were found in nearly all Balderarchaeota MAGs (Fig. 4, Supplementary Table 6). Although the *pta* gene was also found in Hoder-, Sigyn- and Freyrarchaeota, all of their MAGs lack the *ack* gene. The Pta/Ack pathway for acetate production which is common in bacteria but was so far only found in Bathyarchaeota and the methanogenic genus *Methanosarcina* in archaea [43,44] and it is the first time that genes coding for Pta and Ack were discovered in Asgard archaea. The archaeal *pta*/*ack* genes were considered HGT from bacteria donors. For example, the genes in *Methanosarcina* were postulated to acquire from a cellulolytic Clostridia group [45] whereas the *pta*/*ack* genes donor of Bathyarchaeota was still unclear, possibly one unknown clade of Bacteria [43]. For Balderarchaeota, the phylogenetic analysis of the *ack* gene sequences revealed that *ack* genes of Balderarchaeotal branch close to a bacteria lineage Petrotoga (Supplementary Fig. 7), indicating that *ack* genes of Balderarchaeota probably acquired from Petrotoga. While the phylogenetic tree of *pta* genes shows that the Balderarchaeota clade are within Firmicutes branch (Supplementary Fig. 8). Taken together, the Pta/Ack pathway in Balderarchaeota may have acquired from different bacterial donor by two separate HGT events.

The newly discovered Asgard archaea contain less sulfur and nitrogen metabolisms genes (Fig. 4) compared with the archaea in their sister-lineage TACK superphylum [46]. With regard to sulfur metabolism, most of the key enzymes functioning in the assimilatory sulfate reduction pathway and sulfate import were identified in all MAGs of the five new Asgard clades, suggesting that these clades can assimilate sulfate. Moreover, complete subunits of sulfhydrogenase-encoding genes (*hyd*ABGD) were found in all new Asgard lineages except for Balderarchaeota. This bifunctional hydrogenase has been verified in the hyperthermophilic *Pyrococcus furiosus*, which can either remove reductants produced during fermentation by utilizing protons or use polysulfides as electron acceptors [47]. With regard to nitrogen metabolism, only Freyrarchaeota MAGs were found to contain genes coding for the potential nitrogen fixation-catalyzing subunit of nitrogenase (*nif*H) and nitrogenase cofactors. Additionally, genes encoding the nitrite reductase (NADH) large subunit (*nirB*) were detected in MAGs of Sigynarchaeota, implying a potential nitrite reduction capability.

## Conclusion

Undoubtedly, the origin of eukaryotes is one of the most important evolutionary events. The discovery of Asgard archaea has boosted the eocyte hypothesis that eukaryotes derive from within archaea because the Asgard archaea possess two remarkable features: robust evolutionary affinity with eukaryotes and various ESPs existence. In the present study, five novel Asgard lineages were discovered based on phylogenetic analyses and AAI value comparison, which significantly expand the phylogenetic and metabolic diversity of the Asgard archaea. Our analyses strongly support a 2-domain tree of life and clearly demonstrate that the eukaryotes lineage cluster with Tyra-, Heimdall- and Gerdarchaeota, while Tyrarchaeota lineage is a deeper branching lineage to eukaryotes than Heimdallarchaeota. Metabolic characteristic of Tyrarchaeota shows different carbon metabolic pathways from Heimdallarchaeota and Gerdarchaeota that were considered living in aerobic environment using various organic substrates [11,16,48], whereas Tyrarchaeota lack both complete glycolysis and TCA pathway but have potential to utilize hydrogen and WL pathway for carbon fixation. This finding does not contradict to the hypothesis by Spang *et al.* inferring metabolic feature of the archaeal ancestor of eukaryotes [11] because Tyrarchaeota may have closer interaction with their potential bacteria partner as they also contain acetogenesis pathway and the potential ability to synthesize cobalamin, which might be beneficial for symbiosis. In general, the characterization of additional genomes and continuous efforts to cultivate Asgard archaea will provide additional insights into the evolution of archaea and their potential evolution into eukaryotes. Such insights will enable greater understanding of the ecological and geochemical roles of archaea in Earth’s history.

## Methods

The raw data were downloaded from the SRA database and subsequently trimmed, assembled, binned and classified. The metabolic reconstruction, ESPs identification and phylogenetic analyses were conducted. The detail methods were described in Supplementary materials.

## Acknowledgments

We are grateful for Dr. Tom A. William for his suggestions regarding phylogenetic analysis. We thank the researchers who published their metagenomic data on the NCBI website (https://www.ncbi.nlm.nih.gov/), and the datasets used in the current study along with the contributors’ names are listed in Supplemental Table 1.

## Data availability

The genomes of Asgard archaea generated in this study have been made available at the eLibrary of Microbial Systematics and Genomics (eLMSG; https://www.biosino.org/elmsg/index) under accession numbers LMSG_G000000610.1-LMSG_G000000628.1.

## Author contributions

R.Z.X., Y.Z.W. and F.P.W. conceived the study. R.Z.X., Y.Z.W., D.Y.H., H.J.L., H.N.H., L.Y.L. and X.X.Z. analyzed the data. R.Z.X., Y.Z.W. and F.P.W. wrote the paper.

## Compliance and ethics

The author(s) declare that they have no conflicts of interest.

