## Supplemental materials for "Expanded Asgard archaea shed new light on the origin of eukaryotes and support a 2-domain tree of life"

### Supplementary Methods

**Data collection.** Asgard archaea are distributed mainly in different environmental sediments, including estuary sediments [1,2], mangrove sediments [3], hydrothermal sediments [4], hot spring sediments [5], marine sediments [6], and freshwater sediments [7]. According to the environmental distributions of Asgard archaea, metagenomic data were collected and downloaded from the SRA database (https://www.ncbi.nlm.nih.gov/sra/).

**Metagenomic assembly and genomic binning.** The raw reads were trimmed using Trimmomatic (v.0.38) [8] to remove adapters and low-quality reads. After trimming, the reads of each sample were *de novo* assembled using Megahit (v.1.2.5) [9] with a k-step of 6. Samples from the same location or similar environments were assembled together. Contigs were binned separately using MetaBAT (v.2.12.1) [10], MaxBin (v.2.2.7) [11], and Concoct (v.1.1.0) [12] with the default parameters, and the initial taxonomic classification of each MAGs was performed using GTDB-Tk (v.1.2.0) [13] to extract Asgard MAGs. The completeness and contamination of Asgard MAGs were evaluated with the CheckM lineage_wf workflow (v.1.0.12) [14]. Finally, Asgard MAGs with completeness above 50% and contamination below 10% were selected for further analyses, and Prodigal (v.2.6.1) [15] was used to predict protein-coding genes for these selected Asgard MAGs.

**Phylogenetic analyses of Asgard MAGs.** To determine the exact phylogenetic affiliations in the Asgard superphylum, 37 conserved marker genes were selected as described in literatures [16,17]. Homologs of the 37 conserved marker proteins were identified using Diamond (v.2.0.4) [18]. Each dataset of marker proteins was aligned with MAFFT-L-INS-i (v.7.313) [19] and trimmed by trimAl (v.1.4.22) [20] with the "automated1" option.

Maximum-likelihood phylogenies for the 37 conserved marker proteins was built using IQ-TREE (v.2.0.5) [21] under the best-fit model "LG+R10". The support values were calculated using 1000 ultrafast bootstraps.

**Phylogenetic tree of life.** To confirm the phylogenetic affiliations of eukaryotes and the novel Asgard lineages, 21 taxonomic marker genes shared among three domains selected by Williams [22] and 55 ribosomal proteins shared between archaea and eukaryotes [5] were used for phylogenetic analyses. Single-gene trees were inferred for all the markers using IQ-TREE with the LG+G4+F model to exclude eukaryotic genes of mitochondrial or chloroplastic origin or acquired by HGT (if eukaryotic genes which fall into bacterial clade were considered as mitochondrial or chloroplastic origin and that scatter in archaeal clade were treated as HGT) and a BLASTp inspection was further performed to identify all eukaryotic genes originating from the nuclear genome. The maximum-likelihood tree was built with IQ-TREE under the LG+C60+F+G4 model with 1000 ultrafast bootstraps. For Bayesian analysis, four chains were run simultaneously using PhyloBayes-MPI (v.1.8) [23] with the CAT+GTR+G4 model. Four independent chains were run for ~30,000 generations. After a burn-in half of the generations, maxdiff values below 0.3 was achieved.

**Identification of ESPs.** All predicted proteins encoded by the MAGs of the five novel Asgard lineages were analyzed using InterProScan [24] (v.5.47-82.0) with default parameters to annotate protein domains and were assigned to archaeal clusters of orthologous genes (arCOGs) [25] by eggnog-mapper (v. 2.0.1b) [26] with default settings. The lists of InterPro accession numbers (IPRs) and arCOG identifiers previously published by Zaremba-Niedzwiedzka *et al.* [5] and Bulzu *et al.* [27] were used to identify potential ESPs.

Some key words related to eukaryote-specific processes or cell structures were used to search annotation information of interProScan to identify potential ESPs previously not reported in Asgard archaea. Several candidate ESPs were further examined using HHpred [28] with default parameters.

**Metabolic reconstruction.** The proteome of each MAG of the six novel Asgard phyla was uploaded to the KEGG Automatic Annotation Server (KAAS) [29] and run with several settings: the GHOSTX, Prokaryotes and Bidirectional Best Hit (BBH) settings. Additionally, proteins were queried against the nonredundant (NR) protein database (downloaded from NCBI on February 2020) using the Diamond (v.2.0.4) BLASTp search (e-value cutoff $<1e{-}5$). Metabolic pathways were reconstructed based on combination of the NR annotations, protein domain information and KEGG Ontology (KO) numbers.

The dbCAN2[30] web server was used to identify carbohydrate-degrading enzymes with the default settings, and the putative large subunits of [NiFe] hydrogenases were identified by querying against a local database based on HydDB [31] using Diamond (v.2.0.4) with an E-value cutoff of $1\times10^{-20}$. Additionally, a local MEROPS database (downloaded September 2020) [32] searched for peptidases by Diamond (v.2.0.4) with an E-value cutoff of $1\times10^{-20}$, and PSORT (v.3.0.2) was used to identify protein localization [33].

**Calculation of ANI and average AAI.** The ANI and AAI values were calculated using OrthoANI (v.1.2) [34] and CompareM (https://github.com/dparks1134/CompareM), respectively, with the default parameters.

27. Bulzu P-A, Andrei A-Ş, Salcher MM, *et al.* Casting light on Asgardarchaeota metabolism in a

sunlit microoxic niche. *Nature Microbiology* 2019;**4**(7):1129-1137.

### Supplementary Figures

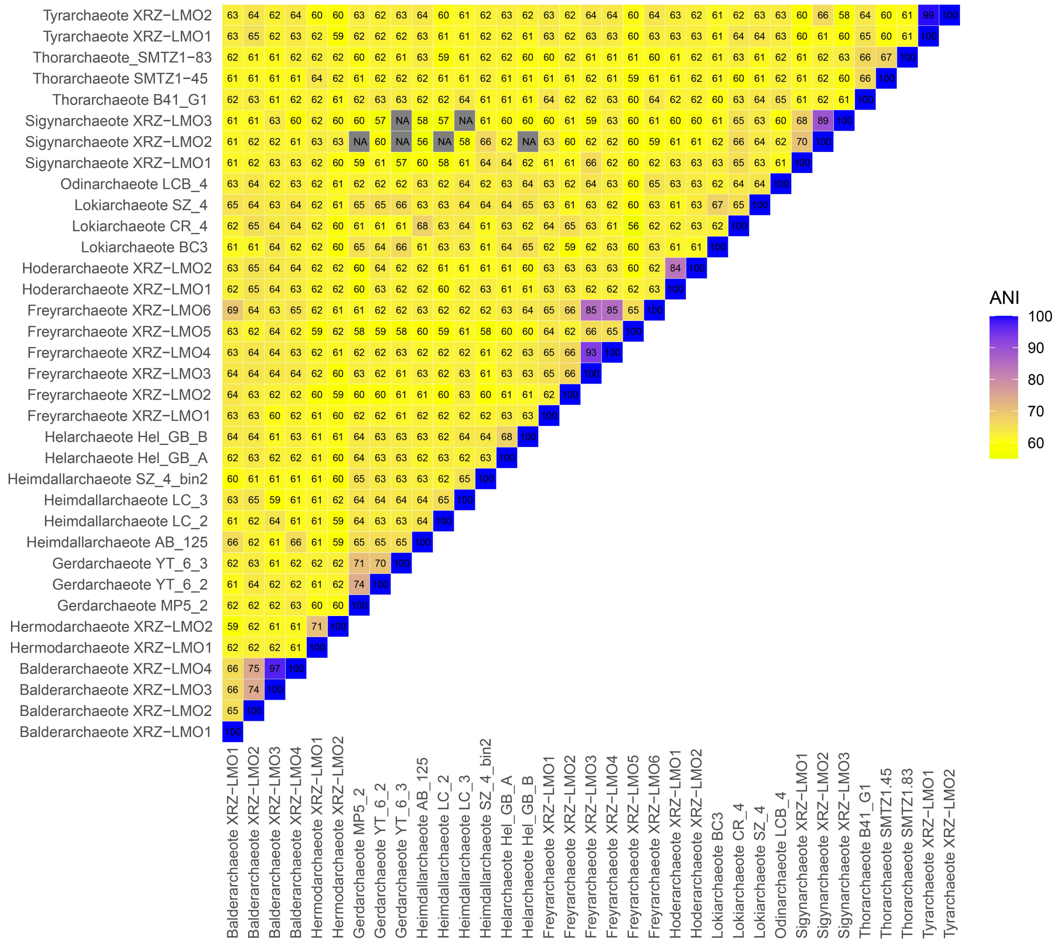


**Supplementary Figure 1.** Average nucleotide identity (ANI) values of the six novel Asgard archaea with the known Asgard archaea as calculated by OrthoANI.

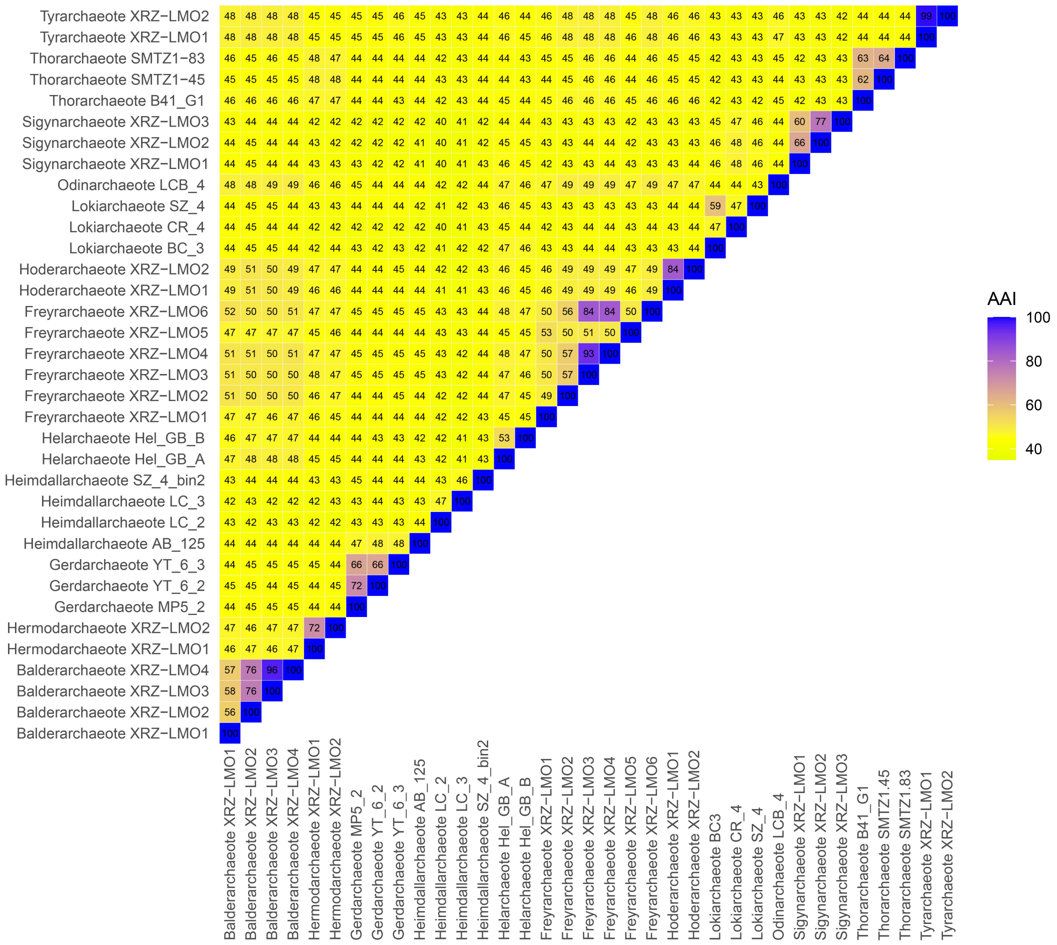


**Supplementary Figure 2.** Amino acid identity (AAI) values of the six novel Asgard archaea with the known Asgard archaea as calculated by CompareM.

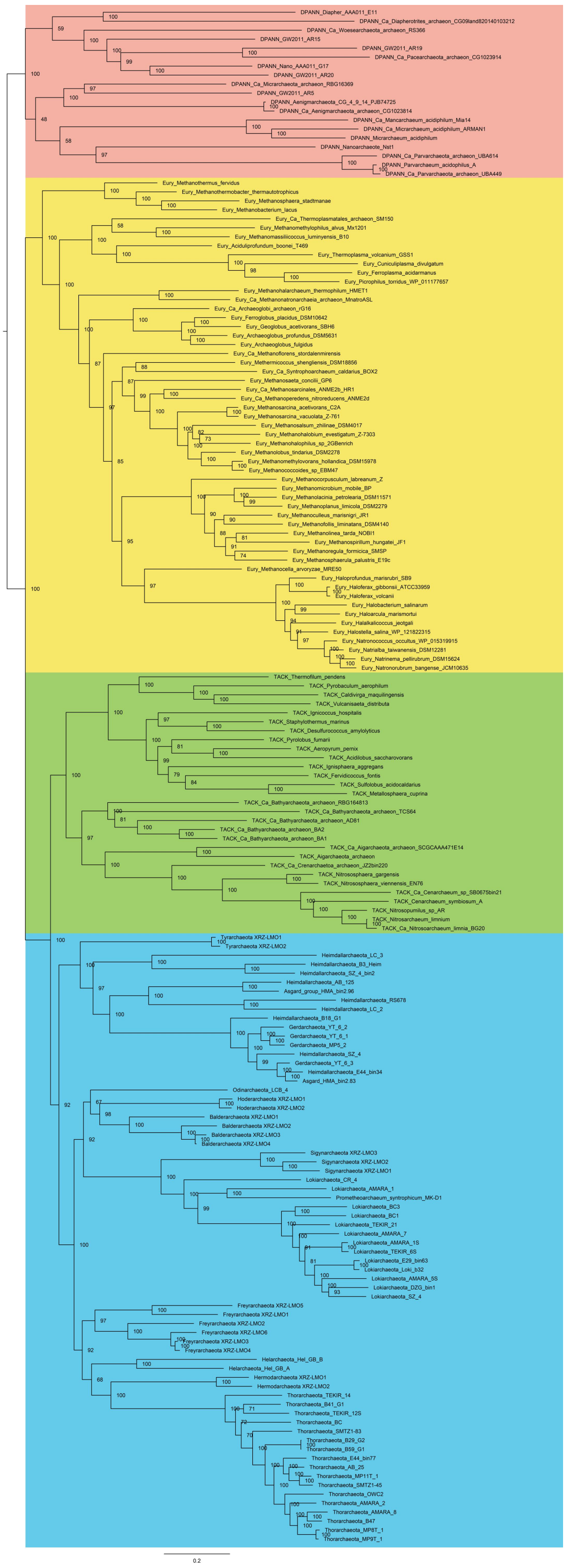

**Supplementary Figure 3** The initial maximum likelihood tree of 37 conserved marker genes under LG+R10 model. DPANN in lightpink, Euryarchaeota in yellow, TACK in green, Asgard in blue.

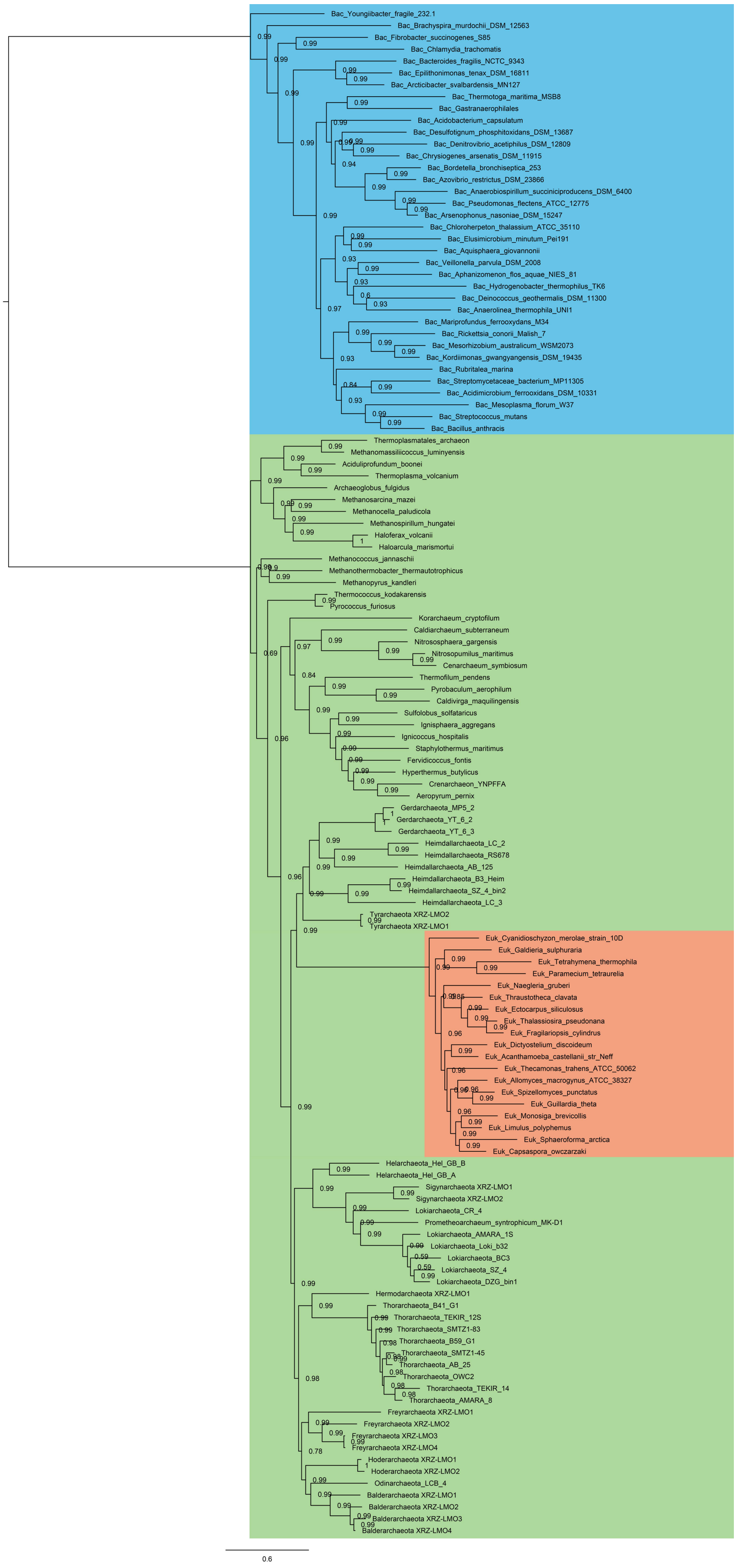


**Supplementary Figure 4** The initial Bayesian tree of 21 conserved marker genes under CAT+GTR+G4 model. Bacteria in blue, archaea in green, eukaryotes in lightsalmon.

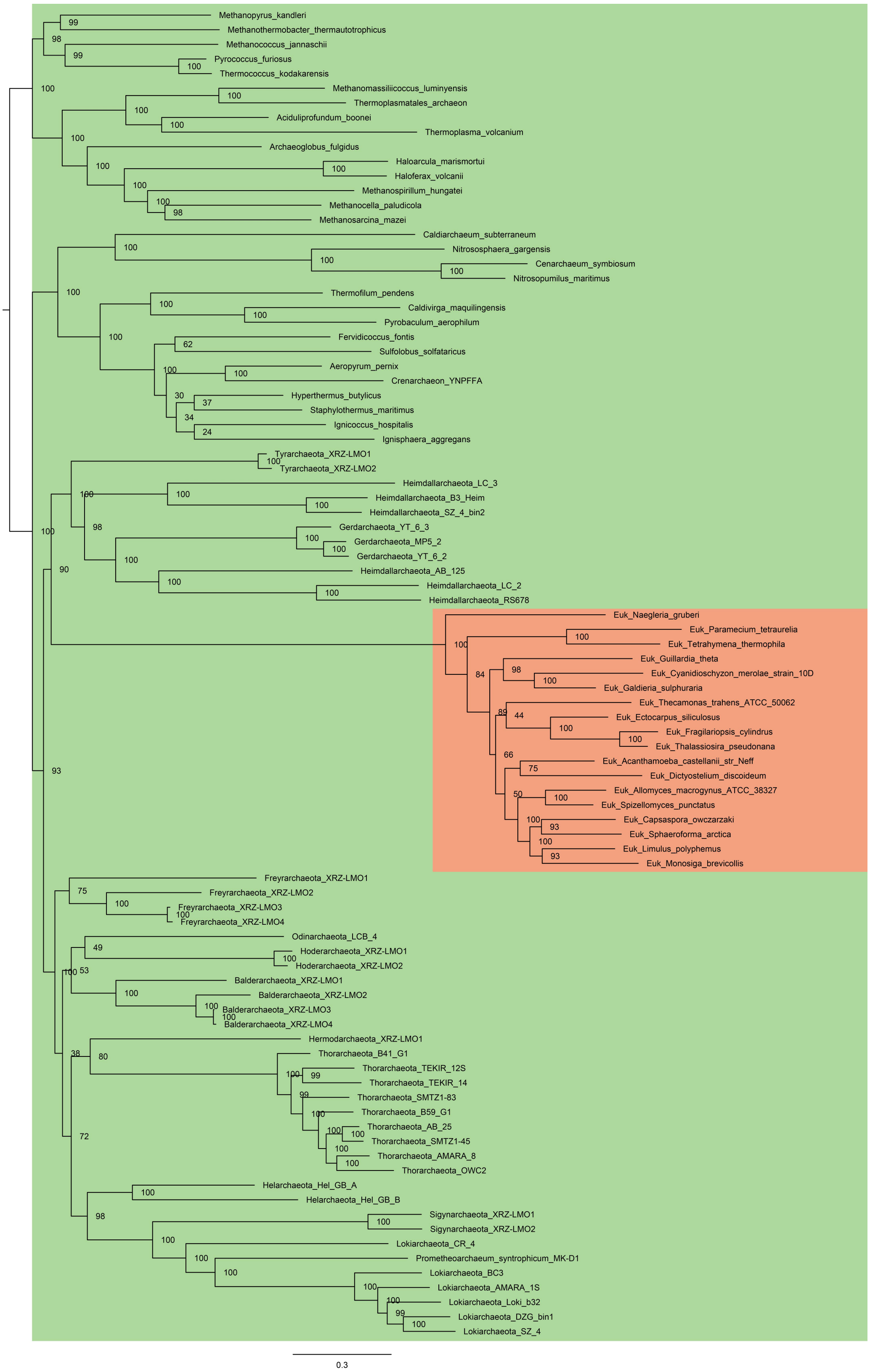


**Supplementary Figure 5** The initial maximum likelihood tree of 55 archaeal-eukaryotic ribosomal proteins under LG+C60+F+G4 model. Archaea are indicated with green; eukaryotes are indicated with lightsalmon.

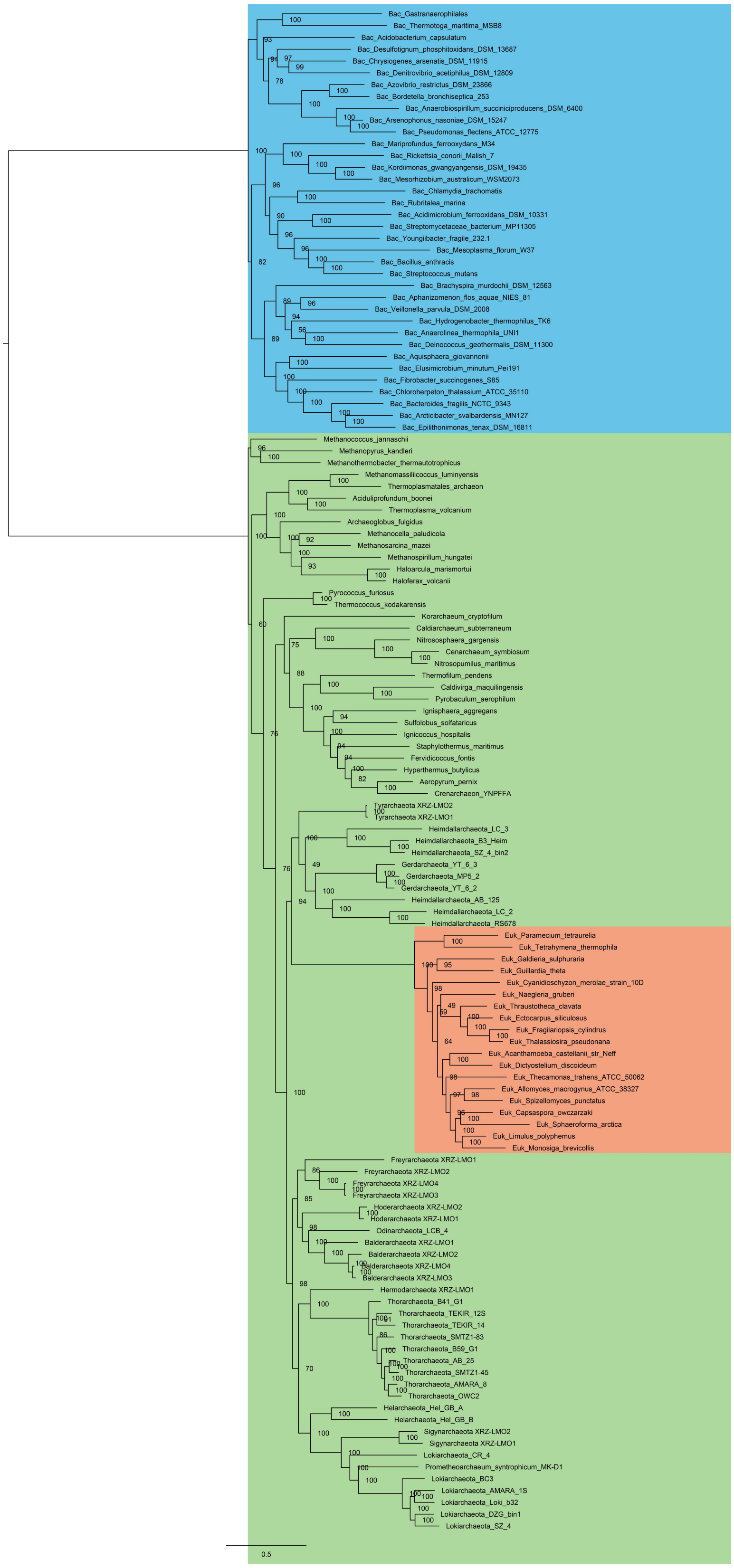


**Supplementary Figure 6** The initial maximum likelihood tree of 21 conserved marker genes after removing inter-archaea HGT under LG+C60+F+G4 model. Bacteria in blue, archaea in green, eukaryotes in lightsalmon.

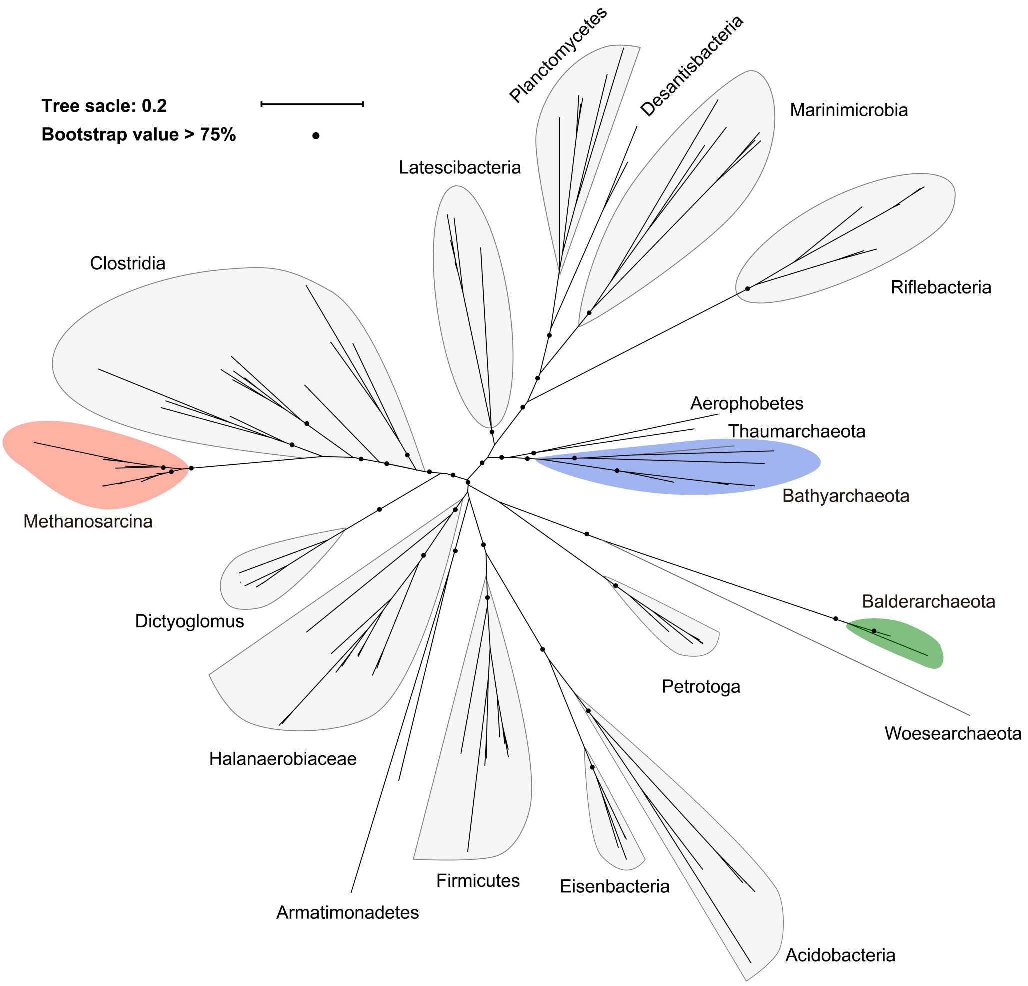


**Supplementary Figure 7** An unrooted maximum-likelihood tree of the Ack homologs using IQ-TREE with best fit model LG+I+G4. Bathyarchaeota in blue, Balderarchaeota in green, Methanosarcina in lightsalmon. The Ack homologs were also found in Woesearchaeota and Thaumarchaeota. Woesearchaeota form a clade with Balderarchaeota and Thaumarchaeota are within Bathyarchaeota lineage, indicating that the *ack* gene in Balderarchaeota and Woesearchaeota acquired from a common bacteria lineage and the *ack* gene in Bathyarchaeota and Thaumarchaeota transferred from a common bacteria donor.

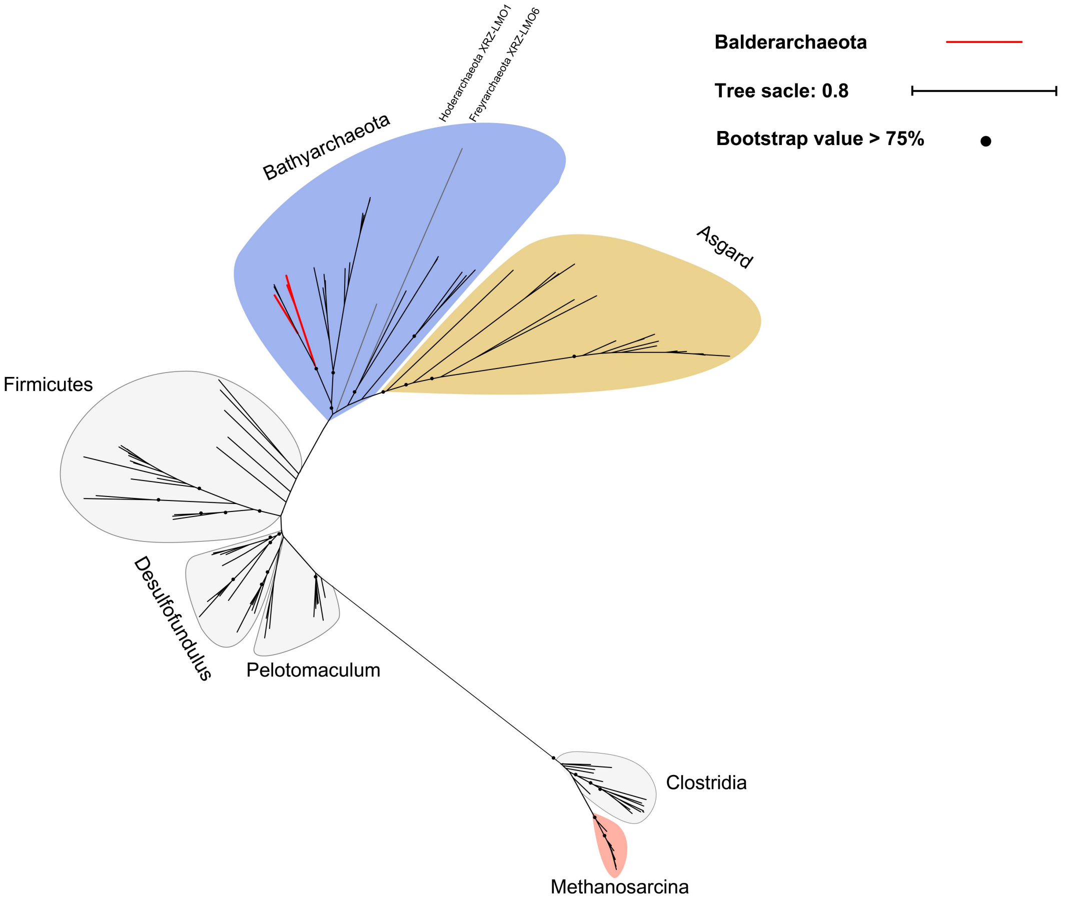


**Supplementary Figure 8** An unrooted maximum-likelihood tree of the Pta homologs using IQ-TREE with best fit model LG+I+G4. Bathyarchaeota in blue, Asgard archaea (except for Balderarchaeota, Hoderarchaeota and Freyrarchaeota) in orange, Methanosarcina in lightsalmon. The red branches are sequences from Balderarchaeota. Both Bathyarchaeota lineage and Asgard lineage are within Firmicutes.
